# BIN1 deficiency leads to DNA damage and neuronal insulin resistance through ATM dysregulation

**DOI:** 10.1101/2025.08.20.671271

**Authors:** Yuxi Jin, Huili Shi, Lin Zhao, Guannan Zhang, Shanshan Ding, Yu Wu, Wen Fan, Zhenghui Liu, Hang Ruan, Yanxiu Li, Ming Xiao, Chengyu Sheng

**Author notes:** These authors contributed equally: Yuxi Jin, Huili Shi, Lin Zhao, Guannan Zhang. Corresponding authors: Chengyu Sheng, Ming Xiao, or Guannan Zhang. Abbreviations: AAV: adeno-associated virus; Aβ: amyloid-β; ACTB: β actin; AD: Alzheimer’s disease; AKT1: thymoma viral proto-oncogene 1; BIN1: bridging integrator 1; BIN1-iso1: BIN1, isoform 1; CA1: cornu Ammonis 1; DMEM: Dulbecco’s modified Eagle medium; GWAS: genome-wide association study; MTOR: mechanistic target of rapamycin kinase; mTORC1: MTOR complex 1; RPS6KB1: ribosomal protein S6 kinase B1; qRT-PCR: real-time quantitative reverse transcription PCR; ROS: reactive oxygen species.

## Abstract

The major neuronal isoform of the AD risk gene BIN1 is specifically reduced in patients. In the present work, we demonstrate that BIN1 is necessary for intact insulin signaling and its loss leads to cellular-level insulin resistance. Persistently activated mTORC1 feedback inhibits the insulin signaling; meanwhile, ATM-dependent DNA Damage Response decreases the expression of multiple insulin signaling pathway members. Unexpectedly, nuclear ATM activation is accompanied by autophagic degradation of the cytoplasmic ATM, rendering neurons susceptible to oxidative stress, which predisposes oxidative DNA damage, DNA strand breaks, and ATM-dependent DNA Damage Response. Treating BIN1-deficient mice with rapamycin or liraglutide to improve the insulin response, or with the natural antioxidant alpha-lipoic acid to attenuate oxidative stress, both effectively preserved the spatial cognitive ability. Lastly, we reanalyzed a single-cell transcriptomic dataset from human neurons and identified positive correlations between levels of BIN1 and the insulin signaling, as well as neuronal activity. Together, this work revealed an ATM-centered mechanism that globally damages neuronal health in BIN1-deficient pyramidal neurons, which could be a driving force in the AD disease course.

## Introduction

BIN1 is a multifunctional protein in both peripheral tissues and the central nervous system (CNS). For late-onset Alzheimer’s disease (AD), genome-wide association studies (GWAS) identified Bridging Integrator 1 (*BIN1*) as the second most significant risk gene.^1, 2^ In the CNS, BIN1 is broadly expressed in neurons, oligodendrocytes, and microglia. The major neuronal isoform of BIN1, BIN1 isoform 1 (BIN1-iso1), is significantly reduced in sporadic AD patients,^3–5^, additionally supporting its participation in the disease course. The downstream consequence of this molecular event is unclear. Clinical studies revealed a close relationship between BIN1 and the microtubule-associated protein TAU.^6–8^ BIN1 directly interacts with TAU,^9–11^ and regulates intercellular propagation of TAU pathology;^12^ overexpression of BIN1 induces neuronal network hyperexcitability in a TAU-dependent manner.^13^ Meanwhile, BIN1 also modulates neurodegeneration independently of TAU. Researchers found that loss of *BIN1* impairs glutamate release at the presynapse and disturbs the membrane targeting of the AMPA receptor at the postsynapse.^14, 15^ Employing the Drosophila larvae model, we previously reported that the ortholog of human *BIN1*, *Amphiphysin*, regulates dendritic branch dynamics and is required for synapse maturation.^16^ Besides these synaptic functions, *BIN1* genotypes were associated with the hippocampal volume in young healthy Chinese university students.^17^ In agreement with this clinical finding, we recently reported that BIN1 deficiency leads to decreased dendritic arbor size in cultured hippocampal neurons with regional volume loss that is visible in magnetic resonance imaging examination of mouse hippocampal CA1; we also proved that elevation of ULK3-initiated, the mammalian target of rapamycin complex 1 (mTORC1)-insensitive autophagic flux is beneath this dendritic atrophy.^18^

In BIN1-deficient neurons, decreased AKT activity facilitates the nuclear translocation of TFEB, which transcriptionally expresses autophagic and lysosomal genes.^18^ As IRS1– PI3K–AKT signaling is a main pathway downstream of insulin binding to its receptor, the reduction of AKT activity suggests the existence of neuronal insulin resistance, which is a common pathology in multiple neurodegenerative disorders, including AD.^19^ Brain insulin resistance typically takes place without systemic insulin resistance in AD patients, and thus, some researchers proposed AD as a type 3 diabetes.^20^ Neuronal insulin resistance is firmly related to cognitive decline through multiple mechanisms.^21^ In the context of AD, Aβ oligomers are known to produce insulin resistance through competing with the insulin receptor, and activated glia-derived pro-inflammatory cytokines such as tumor necrosis factor-α also inhibit IRS1 function through the JNK signaling.^22^ As GSK-3β is negatively regulated by AKT, tau hyperphosphorylation is generally regarded as a downstream event of neuronal insulin resistance. On the risk gene side, ApoE4 promotes the development of insulin resistance through increasing Aβ burden;^23^ whether other risk genes are also linked to neuronal insulin resistance is insufficiently studied.

Unlike AD, patients with another neurodegenerative disorder, ataxia telangiectasia (AT), are prone to develop systemic insulin resistance.^24, 25^ AT is caused by loss-of-function mutations of the gene Ataxia telangiectasia mutated (*ATM*). As a phosphatidylinositol 3-kinase-like kinase (PIKK), ATM regulates diverse metabolic signaling pathways, and ATM deficiency was firmly associated with impaired insulin signaling.^26, 27^ In the nucleus, when DNA double-strand DNA breaks (DSBs) are detected, the Mre11-Rad50-Nbs1 (MRN) complex recruits ATM and leads to its auto-activation; ATM then phosphorylates the histone H2AX to generate a docking platform (γH2AX foci) for the subsequent repair machinery, which is a critical step in DNA damage response (DDR). DDR contains extensive transcriptional and post-transcriptional regulations and affects a number of cellular processes. In a post-mortem study on the frozen frontal cortex of AD patients, high DDR levels were associated with downregulation of multiple important insulin signaling pathway (ISP) members,^28^ supporting a correlation between these two cellular states, but whether DDR is a driver of cellular-level insulin resistance is unclear.

In human fibroblasts and cancer cell lines, loss of BIN1 was associated with activation of ATM in an E2F1-dependent manner.^29^ In the present work, we report that ATM activation and ATM-mediated DDR also occur in BIN1-deficient neurons. Interestingly, we found that this DDR is accompanied by a reduction of cytoplasmic ATM, and both of them impair insulin signaling through distinct routes. What’s more, cytoplasmic ATM reduction renders neurons hypersensitive to reactive oxygen species (ROS) stress, which elicits DNA strand breaks in the nucleus and subsequently nuclear ATM-mediated DDR. In mice with hippocampal neuronal *Bin1* specifically reduced, we evidenced declined spatial cognitive performance, which was significantly attenuated by anti-insulin resistance therapy with the mTORC1 inhibitor rapamycin or the glucagon-like peptide-1 receptor agonist liraglutide, or with a natural antioxidant alpha-lipoic acid targeting the upstream ROS-induced DNA damage. Based on these results, we propose that downstream of neuronal BIN1 deficiency, ATM-centered mechanisms produce intertwined insulin resistance, ROS hypersensitivity, and DNA damage.

## Results

### Persistent mTORC1 activation inhibits insulin signaling in BIN1-deficient neurons

In our recent study, we reported that BIN1 deficiency leads to reduced AKT activity and concurrent activation of mTORC1,^18^ implying feedback inhibition of the IRS1–PI3K– AKT signaling by mTORC1 itself and its downstream kinase RPS6KB1, which is a common route to cellular-level insulin resistance. To make clear whether this occurs, we dissected the cortex from neonatal mice and cultured the pyramidal neurons. On day in vitro (DIV) 5, we transduced neurons with a lentivirus expressing a shRNA targeting all isoforms of mouse *Bin1* under the *RNU6* promoter; On DIV11, cells were collected for western blot (WB) analysis; in plko-sh*Bin1* group, we observed a significant reduction in the Ser473 phosphorylation of AKT; for cells receiving insulin treatment before harvest (100 nM for 10 minutes), p-AKT-Ser473 levels increased In the control group, but displayed no significant change in plko-sh*Bin1* group (**Figure 1A**). Meanwhile, *Bin1* knockdown substantially decreased the protein levels of insulin receptor substrate 1 (IRS1) (**Figure 1A**), additionally indicating impaired insulin signaling. To see whether this ISP disturbance is specific to developing neurons, we decreased *Bin1* expression via RNAi in the HT22 hippocampal neuronal cell line. We found that a transient transfection of a sh*Bin1*-expressing plasmid (pGPU6-sh*Bin1*) is sufficient to produce a similar insulin resistance state (**Figure 1B**); though insulin treatment still evoked a significant increase in p-AKT-Ser473 levels in the RNAi group, the magnitude is much smaller (**Figure 1B**).

**Figure 1.**
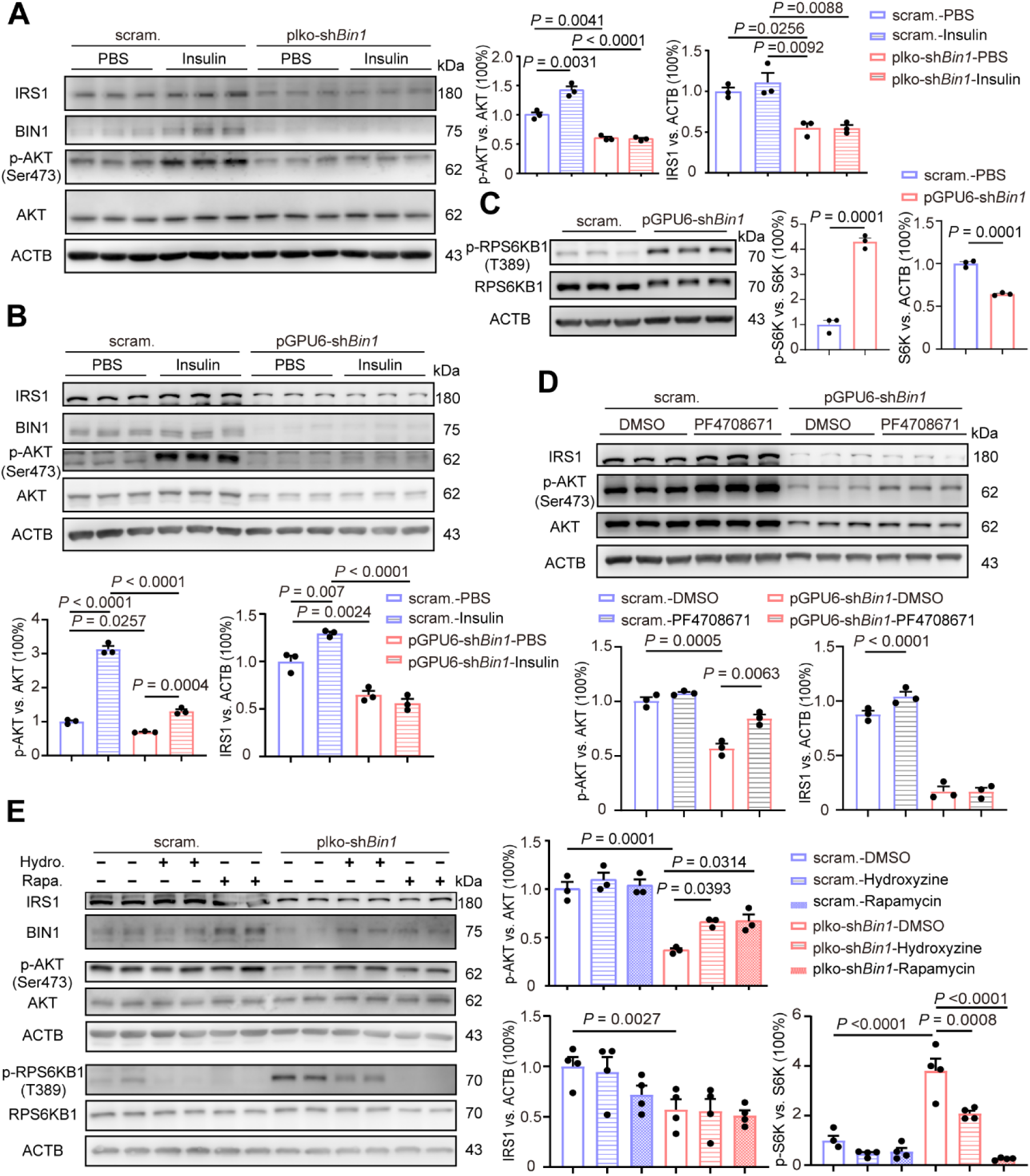
(**A**) Representative WB images showing the expression of IRS1, p-AKT-Ser473, AKT, and BIN1 protein in primary mouse cortex neurons in scramble and plko-sh*Bin1* groups with or without insulin treatment (100 nM, 10 min). Right, a bar graph (Mean + SEM) overlaid with dot plots (individual data points) showing the corresponding quantification of IRS1 and p-AKT-Ser473 bands intensity. n = 3 per group. (**B**) Representative WB images showing the expression of IRS1, p-AKT-Ser473, AKT, and BIN1 protein of HT22 cells in scramble and pGPU6-sh*Bin1* groups with or without insulin treatment (100 nM, 10 min). Right, a bar graph (Mean + SEM) overlaid with dot plots (individual data points) showing the corresponding quantification of IRS1 and p-AKT-Ser473 bands intensity. n = 3 per group. (**C**) Representative WB images and a bar graph (Mean + SEM) overlaid with dot plots (individual data points) showing p-RPS6KB1 and RPS6KB1 protein expression in different groups of HT22 cells. n = 3 per group. (**D**) Representative WB images showing protein levels of IRS1, p-AKT-Ser473, and AKT in scramble and pGPU6-sh*Bin1* groups with or without PF4708671 treatment (10 μM, 1 h); n = 3 per group. (**E**) Representative WB images showing protein levels of IRS1, p-AKT-Ser473, AKT, p-RPS6KB1, RPS6KB1, and BIN1 in scramble and plko-sh*Bin1* groups with or without hydroxyzine (10 μM, 2 h) and rapamycin (200 nM, 2 h) treatment; n = 4 per group.

As in primary neurons, *Bin1* knockdown leads to increased mTORC1 activity in HT22 cells, as indicated by the significantly elevated p-RPS6KB1-Tyr389 levels (**Figure 1C**). It is known that both mTORC1 and RPS6KB1 phosphorylate IRS1 at serine residues and attenuate its activity as well as protein stability. We thus treated the cells with PF4708671, a RPS6KB1 inhibitor, and found that the AKT activity was partially restored (**Figure 1D**). Next, we treated the primary cortical neurons with the pan-mTOR inhibitor rapamycin (200 nM, 2 h), or a mTORC1-specific inhibitor hydroxyzine (10 μM, 2 h),^30^ and checked the ISP activity. As expected, two hours of rapamycin/hydroxyzine treatment effectively reduced p-RPS6KB1-Tyr389 levels and successfully increased AKT activity (**Figure 1E**). Taken together, these results support that the persistent activated mTORC1 and RPS6KB1 feedback inhibits the upstream IRS1–PI3K–AKT signaling, operating as a route of insulin resistance downstream of BIN1 deficiency.

### Rapamycin and liraglutide improve spatial cognitive performance in AAV-shBin1 mice

Neuronal insulin signaling is an important regulator of synaptic plasticity, and its state is tightly associated with cognitive functions.^31^ As brain insulin resistance is common to AD patients, antidiabetic drugs are being investigated to be repurposed for AD treatment. Among them, the Glucagon-like peptide-1 (GLP-1) receptor agonist liraglutide is capable of crossing the BBB and attenuating insulin resistance both directly and indirectly.^32, 33^ Therefore, we explored whether liraglutide and the mTOR inhibitor rapamycin could attenuate insulin resistance and improve cognitive performance in BIN1-deficient mouse models.

As in our previous study, we engineered an adeno-associated virus (AAV9) carrying the identical *Bin1*-targeting shRNA sequence used in our lentiviral constructs. This AAV also expresses a *mCherry* reporter driven by the *Camk2a* promoter, enabling convenient visualization of successfully transduced excitatory neurons (designated as AAV-*Camk2a*-*mCherry*-sh*Bin1*, hereafter abbreviated as AAV-sh*Bin1*; control groups received AAV expressing scrambled shRNA, denoted as AAV-scram.). With stereotaxic intracranial injection, we precisely delivered the AAVs to the hippocampal CA1 subregion in mice. (**Figure 2A**). One month after the injection, mice received intraperitoneal injections of either liraglutide (200 μg/kg) or rapamycin (2 mg/kg) daily for four weeks. We then employed the Barnes maze test to assess the hippocampus-dependent spatial cognition. Mice were first habituated and spent 5 days in the acquisition training phase, at the end of which all mice could find the escape tunnel at a similar time scale. 3 days after the final training session, mice underwent the probe trial to assess their spatial reference memory. In this session, AAV-sh*Bin1* mice exhibited significantly longer target hole latency and more incorrect explorations than controls. Results showed that liraglutide treatment significantly improved spatial memory performance, as evidenced by reduced latency to locate the target hole and fewer incorrect exploration attempts (**Figure 2B**); similarly, rapamycin treatment also reduced the latency, though it did not reduce the number of incorrect attempts (**Figure 2B**). In agreement with impaired spatial cognitive performance, CA1 slices from AAV-sh*Bin1* mice displayed significantly reduced c-Fos levels (**Figure 2C**), indicating reduced neuronal activity. In HT22 cells, BIN1 deficiency also leads to a decrease in c-Fos protein (**Figure S1**). Analyzing CA1 lysates with WB, we observed significantly increased phosphorylation of eIF4E binding protein 1 (4EBP1) on Thr37/46 in the AAV-sh*Bin1* group (**Figure 2D**), indicating elevated mTORC1 activity; meanwhile, AAV-sh*Bin1* mice displayed significantly reduced IRS1 and p-AKT-Ser473 levels, with the latter partially restored by liraglutide or rapamycin treatment (**Figure 2E**). In summary, these *in vivo* data agree with results from cellular models and support persistent mTORC1 activation as a conserved molecular mechanism driving insulin resistance in BIN1-deficient neurons.

**Figure 2.**
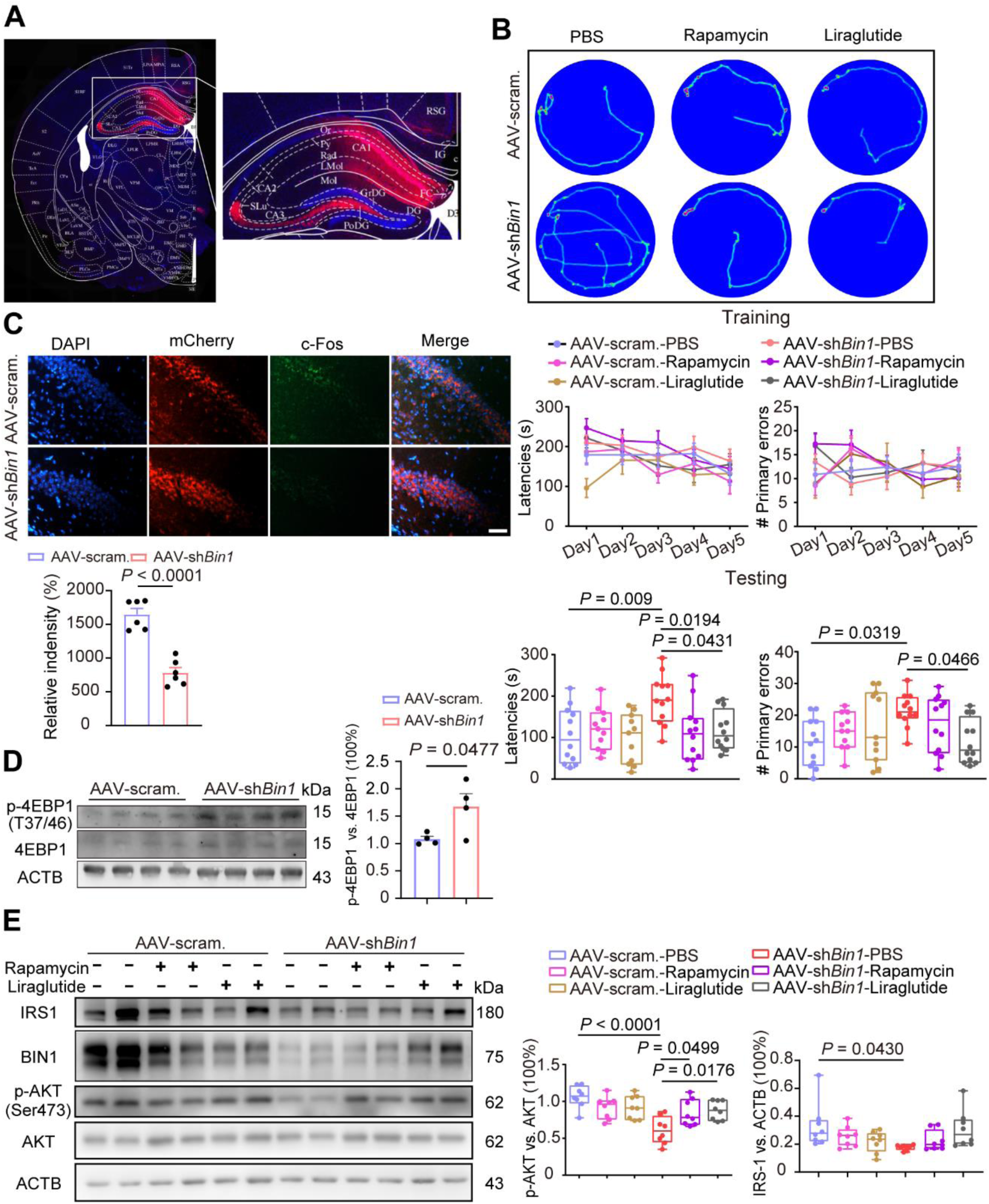
(**A**) An illustration showing stereotaxic intracranial injection into the hippocampal CA1 region of 2-month-old mice. (**B**) Heat maps showing representative trajectories of mice in the Barnes maze during the probe phase; bottom, line graphs showing the latencies and mistakes made (primary errors) before reaching the escape tunnel during the acquisition training phase; bar graphs (Mean + SEM) overlaid with dot plots (individual data points) showing the latencies and primary errors before reaching the previous escape tunnel location during the probe phase. n = 12 mice per group. (**C**) Representative immunofluorescence images showing c-Fos^+^ cells in hippocampal CA1; right, a bar graph (Mean + SEM) overlaid with dot plots (individual data points) showing the numbers of c-Fos^+^ cells in different groups. Scale bar: 100 μm. n = 6 per group. (**D**) Representative WB images and a bar graph (Mean + SEM) overlaid with dot plots (individual data points) showing p-4EBP1(T37/46) and 4EBP1 protein expression in scramble and AAV-sh*Bin1* groups. n = 4 per group. (**E**) Representative WB strips showing protein levels of IRS1, p-AKT-Ser473, AKT, and Bin1 from CA1 lysates in different groups; right, a bar graph (Mean + SEM) overlaid with dot plots (individual data points) showing the corresponding quantification of IRS1, p-AKT-Ser473, AKT, and Bin1 bands intensity. n = 8 per group.

### Reduced TSC1/TSC2 support persistent mTORC1 activation in BIN1-deficient neurons

Why does BIN1 deficiency lead to persistent mTORC1 activation? mTORC1 is directly activated by the small GTPase Ras homolog enriched in brain (RHEB), which is inhibited by the tuberous sclerosis complex 1 (TSC1)–TSC2 complex. In patients with tuberous sclerosis, loss-of-function mutations of the *TSC1* or *TSC2* gene lead to persistent mTORC1 activation. Thus, we examined the protein levels of TSC1 and TSC2 with WB. In cultured cortical neurons, *Bin1* knockdown specifically reduced TSC1 levels (**Figure 3A**); in HT22 cells, *Bin1* knockdown decreased both TSC1 and TSC2, while their phosphorylation (p-TSC1-Ser511 and p-TSC2-Thr1462) remained unchanged (**Figures 3B** and **3C**). At the transcript level, *Bin1* knockdown significantly decreased *TSC1*, while *TSC2* displayed a trend of reduction in the pGPU6-sh*Bin1* group (**Figure 3D**). To investigate whether post-transcriptional mechanisms are also involved, we treated HT22 cells with the translation elongation inhibitor cycloheximide (CHX) (10 μM, 2 h or 4 h) and the proteasome inhibitor MG132 (10 μM, 2 h or 4 h). In the control group, MG132 treatment alone led to a time-dependent accumulation of TSC1 proteins, which was blocked by the additional CHX treatment, indicating that those proteins are newly synthesized; in the pGPU6-sh*Bin1* group, however, no significant TSC1 accumulation was observed with the MG132 treatment alone (**Figure 3E**), suggesting impaired protein translation of TSC1. It thus appears that there are both transcriptional and translational mechanisms are responsible for the dysregulation of TSC1 when BIN1 is deficient.

**Figure 3.**
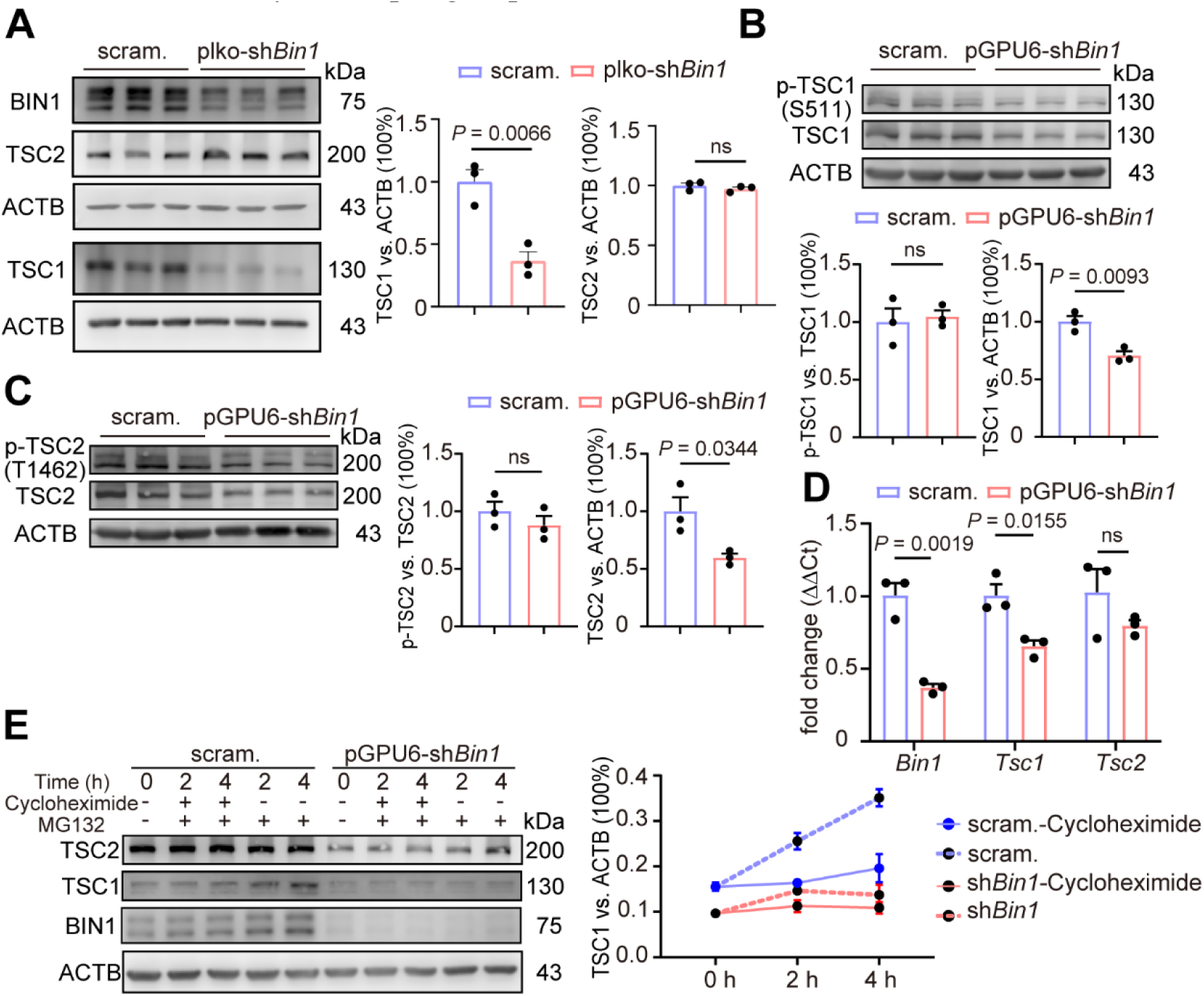
(**A**) Representative WB images and a bar graph (Mean + SEM) overlaid with dot plots (individual data points) showing TSC1, TSC2, and BIN1 protein expression in scramble and plko-sh*Bin1* groups. n = 3 per group. (**B**) Representative WB images and a bar graph (Mean + SEM) overlaid with dot plots (individual data points) showing p-TSC1-Ser511 and TSC1 protein expression in different groups of HT22 cells. n = 3 per group. (**C**) Representative WB images and a bar graph (Mean + SEM) overlaid with dot plots (individual data points) showing p-TSC2-Thr1462 and TSC2 protein expression in different groups of HT22 cells. n = 3 per group. (**D**) A bar graph (Mean + SEM) overlaid with dot plots (individual data points) showing the relative transcript levels of *Bin1, Tsc1,* and *Tsc2* in different groups of HT22 cells. Transcripts were examined by qRT-PCR with *Actb* as an internal reference. n = 3 per group. (**E**) Representative WB images showing the expression of TSC1, TSC2, and BIN1 protein of HT22 cells in different groups with or without cycloheximide (10 μM, 2 h or 4 h) or MG132 (10 μM, 2 h or 4 h) treatment. Right, a line chart 1 (Mean ± SEM) showing the TSC1 protein expression in different groups. n = 3 per group.

### ATM-mediated DNA damage response leads to insulin resistance in BIN1-deficient neurons

Why do TSC1/TSC2 and IRS1 reduce when BIN1 is deficient? In addition, specifically in HT22 cells, total AKT appears to decrease in the RNAi group (**Figures 1B**, **1D**). How could *Bin1* knockdown affect so many ISP proteins? The elevated mTORC1 activity precludes an explanation of global translational repression. What’s more, though treatments with rapamycin or liraglutide attenuated insulin resistance, they primarily affect the phosphorylation of AKT but leave IRS1 levels unaltered (**Figures 1E**, **2E**). Thus, there must be other mechanisms involved in the regulation of the insulin signaling following BIN1 knockdown. We recently reported that BIN1 deficiency induces ULK3-initiated autophagy elevation, which likely degrades some of the ISP members.^34^ However, it turned out that neither attenuating ULK3 activity nor inhibiting lysosomal proteases rescued ISP protein levels (**Figure S2, S3**), demonstrating that the autophagy-lysosome pathway is not responsible.

In human fibroblasts and cancer cell lines, loss of BIN1 leads to ATM activation.^35^ As ATM is an important mediator in DSBs-induced DDR, we hypothesize that BIN1 deficiency also elicits DDR in neurons. DDR activates a cascade of signaling events, including cell cycle arrest, DNA damage repair, and metabolic pathway remodeling, ultimately influencing the global cell state and cellular fate.^36^ Noteworthy, DDR contains transcriptional and post-transcriptional alterations and could potentially alter ISP protein levels. Once activated, ATM phosphorylates H2AX to produce γH2AX foci, serving as a sensitive biomarker for detecting DSBs. In pGPU6-sh*Bin1* cells, we noticed significantly increased γH2AX levels, which were attenuated by co-treating cells with the ATM inhibitor KU60019 (3 μM, 12 hours) (**Figure 4A**), demonstrating that BIN1 deficiency also leads to ATM-mediated DDR in HT22 cells. Interestingly, the KU60019 treatment almost completely restored protein levels of TSC1, TSC2, RPS6KB1, IRS1, and AKT in pGPU6-sh*Bin1* cells (**Figures 4B** and **4C**), and effectively restored cellular insulin response as evaluated by the increase of p-AKT-Ser473 levels following insulin treatment (**Figure 4C**). These dramatic and extensive effects strongly support DDR as a main driver of insulin resistance in BIN1-deficient cells. Similar to HT22 cells, BIN1 deficiency also increased γH2AX levels in primary cortical neurons (**Figure 4D**), and KU60019 treatment dramatically restored ISP protein levels in the plko-sh*Bin1* group, though AKT is an exception (**Figure 4E**). Unlike in proliferating cells, ATM in neurons is predominantly in the cytoplasm, where it acts as a PI3K family member that can directly phosphorylate AKT at Ser 473 to activate it,^37^ which explains why KU60019 did not improve AKT phosphorylation in cortical neurons as in HT22 cells (**Figure 4E**). Collectively, these data support that ATM-mediated DDR is a main driver of neuronal insulin resistance when BIN1 is deficient.

**Figure 4.**
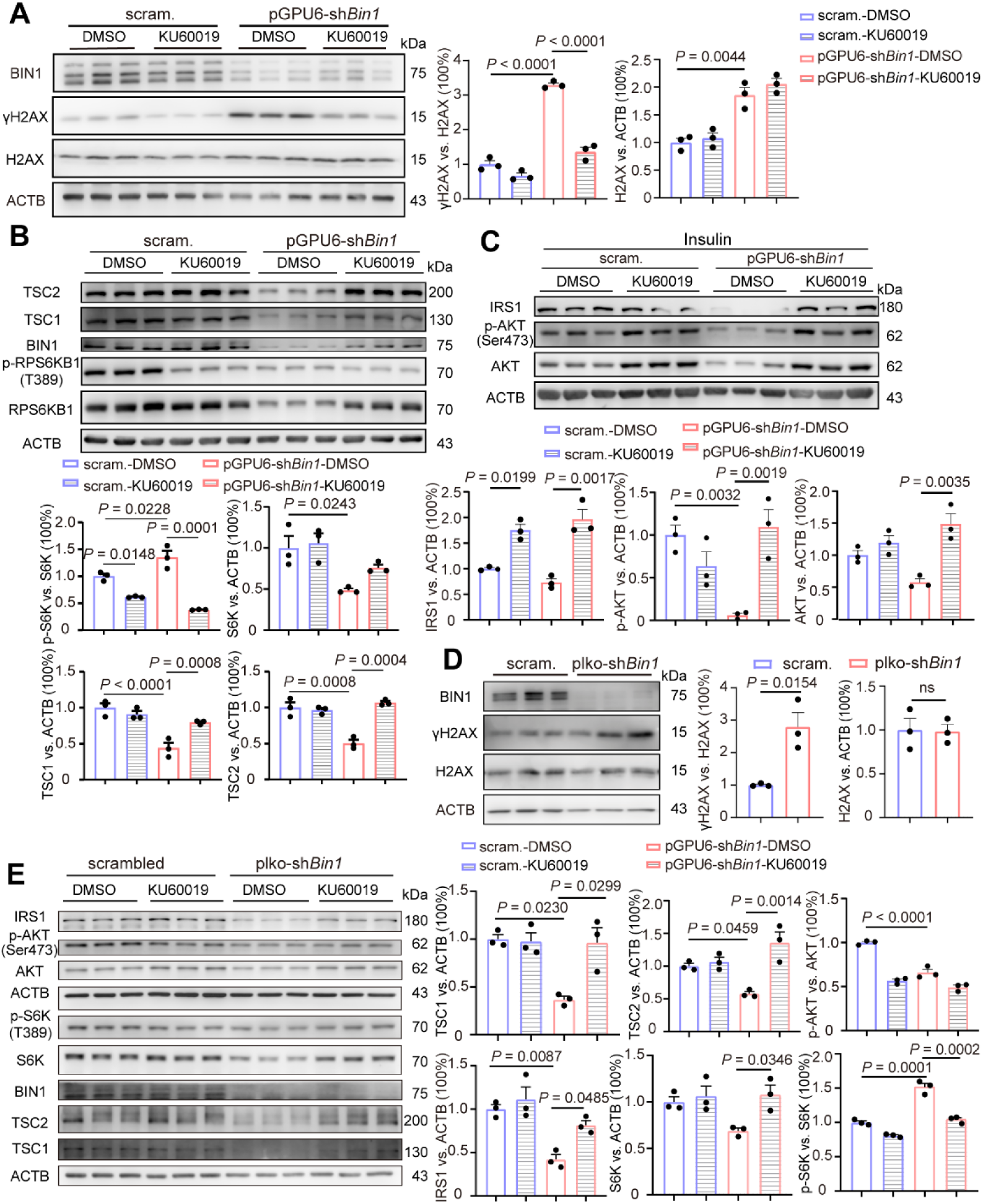
(**A**) Representative WB images showing protein levels of γH2AX, H2AX, and Bin1 in scramble and pGPU6-sh*Bin1* groups with or without KU60019 treatment (3 μM, 72 h); n = 3 per group. (**B**) Representative WB images showing protein levels of p-RPS6KB1, RPS6KB1, TSC1, TSC2, and BIN1 in scramble and pGPU6-sh*Bin1* groups with or without KU60019 treatment (3 μM, 72 h); n = 3 per group. (**C**) Representative WB images showing protein levels of IRS1, p-AKT-Ser473 and AKT in scramble and pGPU6-sh*Bin1* groups with or without KU60019 (3 μM, 72 h) and insulin (100 nM, 10 min) treatment; n = 3 per group. (**D**) Representative WB images and a bar graph (Mean + SEM) overlaid with dot plots (individual data points) showing γH2AX, H2AX and BIN1 protein expression in different groups of primary cortex neurons. n = 3 per group. (**E**) Representative WB images showing protein levels of p-RPS6KB1, RPS6KB1, TSC1, TSC2, IRS1, p-AKT-Ser473, AKT and BIN1 in scramble and plko-sh*Bin1* groups with or without KU60019 treatment (3 μM, 72 h); n = 3 per group.

### BIN1 deficiency decreases ATM protein levels, which also suppresses insulin signaling

Following DSBs are sensed by the MRN complex, ATM is recruited and undergoes self-phosphorylation at Ser1981.^38^ In BIN1-deficient HT22 cells, p-ATM-Ser1981 levels are significantly elevated, confirming the activation of ATM; unexpectedly, ATM protein itself is significantly reduced in pGPU6-sh*Bin1* cells (**Figure 5A**). Since ATM is a PI3K kinase, the change of its protein level provides another layer of regulation for the insulin signaling. Similar to HT22 cells, BIN1 deficiency also leads to a significant reduction of ATM proteins in cultured pyramidal neurons (**Figure 5B**), and the decrease of ATM is also evidenced in hippocampal CA1 lysates extracted from AAV-sh*Bin1* mice (**Figure 5C**). This raises the question of why ATM reduction does not cancel ATM-mediated DDR in BIN1-deficient cells. One explanation is that BIN1 deficiency drives ATM redistribution to the nucleus to support the DDR. We performed immunofluorescence to evaluate ATM distribution in cultured hippocampal neurons; nevertheless, quantitative analysis revealed no significant difference in terms of ATM subcellular distribution between control and plko-sh*Bin1* groups (**Figure 5D**). It thus appears that a small subset of total ATM protein is already sufficient for DDR activation in the nucleus, which operates in parallel with cytoplasmic ATM reduction and together drives the development of insulin resistance.

**Figure 5.**
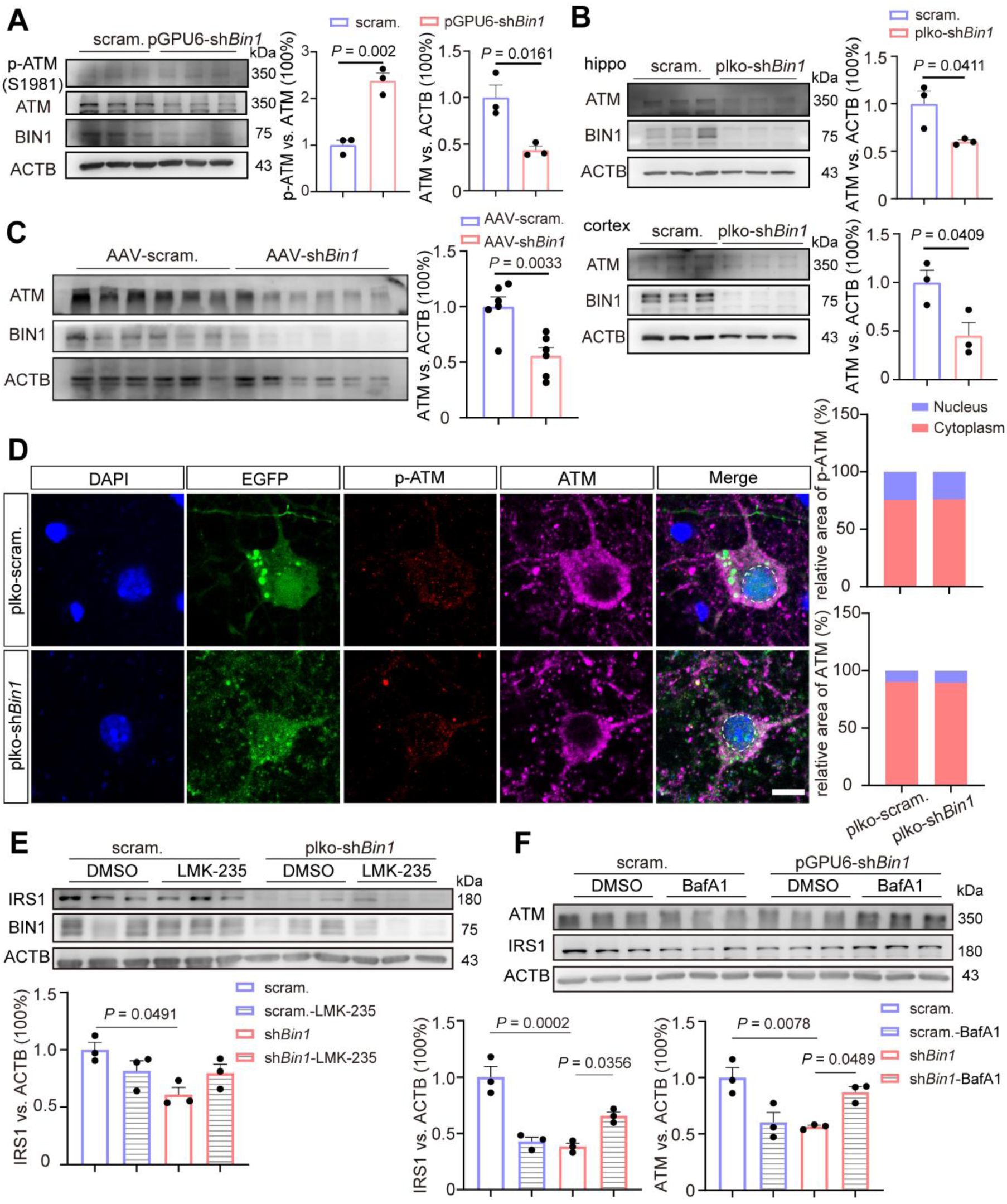
(**A**) Representative WB images showing protein levels of p-ATM and ATM in scramble and pGPU6-sh*Bin1* groups; n = 3 per group. (**B**) Representative WB images and a bar graph (Mean + SEM) overlaid with dot plots (individual data points) showing ATM and BIN1 protein expression in different groups of primary cortex neurons and primary hippocampal neurons. n = 3 per group. (**C**) Representative WB images and a bar graph (Mean + SEM) overlaid with dot plots (individual data points) showing ATM and BIN1 protein expression in the scramble and AAV-sh*Bin1* groups. n = 6 per group. (**D**) Representative immunofluorescence images showing the distribution of p-ATM and ATM of hippocampal neurons in scramble and plko-sh*Bin1* groups; bottom, a bar graph (Mean + SEM) overlaid with dot plots (individual data points) showing the percentage of area occupied by p-ATM and ATM in the nucleus or cytoplasm in different groups; Scale bar: 10 μm. n = 11-12 per group. (**E**) Representative WB images and a bar graph (Mean + SEM) overlaid with dot plots (individual data points) showing IRS-1 and BIN1 protein expression in scramble and plko-sh*Bin1* groups with or without LMK-235 treatment (200 nM, 24 h); n = 3 per group. (**F**) Representative WB images and a bar graph (Mean + SEM) overlaid with dot plots (individual data points) showing IRS-1 and ATM protein expression in scramble and pGPU6-sh*Bin1* groups with or without BafA1 treatment (0.1 μM, 24 h); n = 3 per group.

In the cytoplasm, neuronal ATM deficiency was reported to cause nuclear accumulation of histone deacetylase 4 (HDAC4),^39^ which may lead to insulin resistance through epigenetic silencing of *GLUT4* and possibly also ISP members. ^40^ To make clear whether this route is involved, we treated primary neurons with a Class IIa HDAC-selective inhibitor, LMK-235, and examined IRS1 protein levels with WB. In both control and sh*Bin1* groups, we observed no significant alteration of IRS1 levels with the HDAC4 inhibition (**Figure 5E**), indicating the HDAC4-mediated epigenetic mechanism is not involved in IRS regulation. Next, we asked how ATM protein levels are regulated by BIN1 deficiency. As ATM was degraded through the autophagy pathway,^41^ BIN1-deficiency induced autophagy elevation is a potential mechanism.^18^ Thus, we attenuated lysosome acidification with the vacuolar H^+^-ATPase (V-ATPase) inhibitor Bafilomycin A1 (BafA1) in HT22 cells; as expected, we evidenced significant accumulation of ATM proteins in the sh*Bin1* group (**Figure 5F**), supporting an autophagy route of ATM regulation by BIN1.

### ROS causes DNA damage in BIN1-deficient neurons

In BIN1-deficient human fibroblasts and tumor cells, the activation of ATM is regarded to be E2F1-dependent, but is not caused by genuine DSBs.^29^ DSBs have long been found in AD patients and have recently been identified as an important driver of the disease course,^42, 43^ It is important to determine whether BIN1-deficiency elicits DSBs. Firstly, we employed the comet assay to evaluate whether there are DNA strand breaks in BIN1-deficient cells. In both cultured cortical neurons and HT22 cells, we observed significantly more “comet tails” in the RNAi groups (**Figure 6A**), indicating the existence of true DNA damage. Cytoplasmic ATM is a key ROS sensor that helps to maintain the redox homeostasis; ^26^ with reduced available ATM, BIN1-deficiency could compromise cells’ ability to confront ROS stress. To test this, we treated HT22 cells with a low dose of glutamate (5 μM, 24 h) to elicit a mild excitotoxicity. As expected, we observed a mild increase in the cellular ROS level in the control group; in the RNAi group, however, though their basal ROS level is not significantly different, the glutamate treatment elicited significantly higher ROS levels (**Figure 6B**). It thus appears that BIN1 deficiency impairs cells’ ability to eliminate ROS. Excessive ROS may cause oxidative DNA damage. In agreement with the increased ROS susceptibility, a low concentration of H_2_O_2_ challenge that could be effectively handled by normal HT22 cells significantly elevated the 8-oxo-deoxyguanosine (8-oxo-dG) levels in the RNAi group (**Figure 6C**). Besides oxidative DNA damage, ROS could cause DNA strand breaks and mainly single-strand breaks (SSBs), which may also arise from the repair of the oxidized bases. Since the comet assay is insensitive to discriminate SSBs and DSBs, this raises the question whether BIN1 deficiency-induced DNA damage contains genuine DSBs. We employed immunofluorescence analysis of the γH2AX foci to address this concern. In cultured cortical neurons, we observed significantly more γH2AX foci in the plko-sh*Bin1* group without any additional treatment (**Figure 6D**); treating neurons with the ATM inhibitor KU60019 or an antioxidant N-acetyl-L-cysteine (NAC) both effectively eliminated them (**Figure 6D**). Therefore, BIN1 deficiency does induce γH2AX foci formation in an ATM-dependent and ROS-dependent manner, which is in agreement with a previous study that H_2_O_2_ elicits ATM-dependent formation of γH2AX foci.^44^

**Figure 6.**
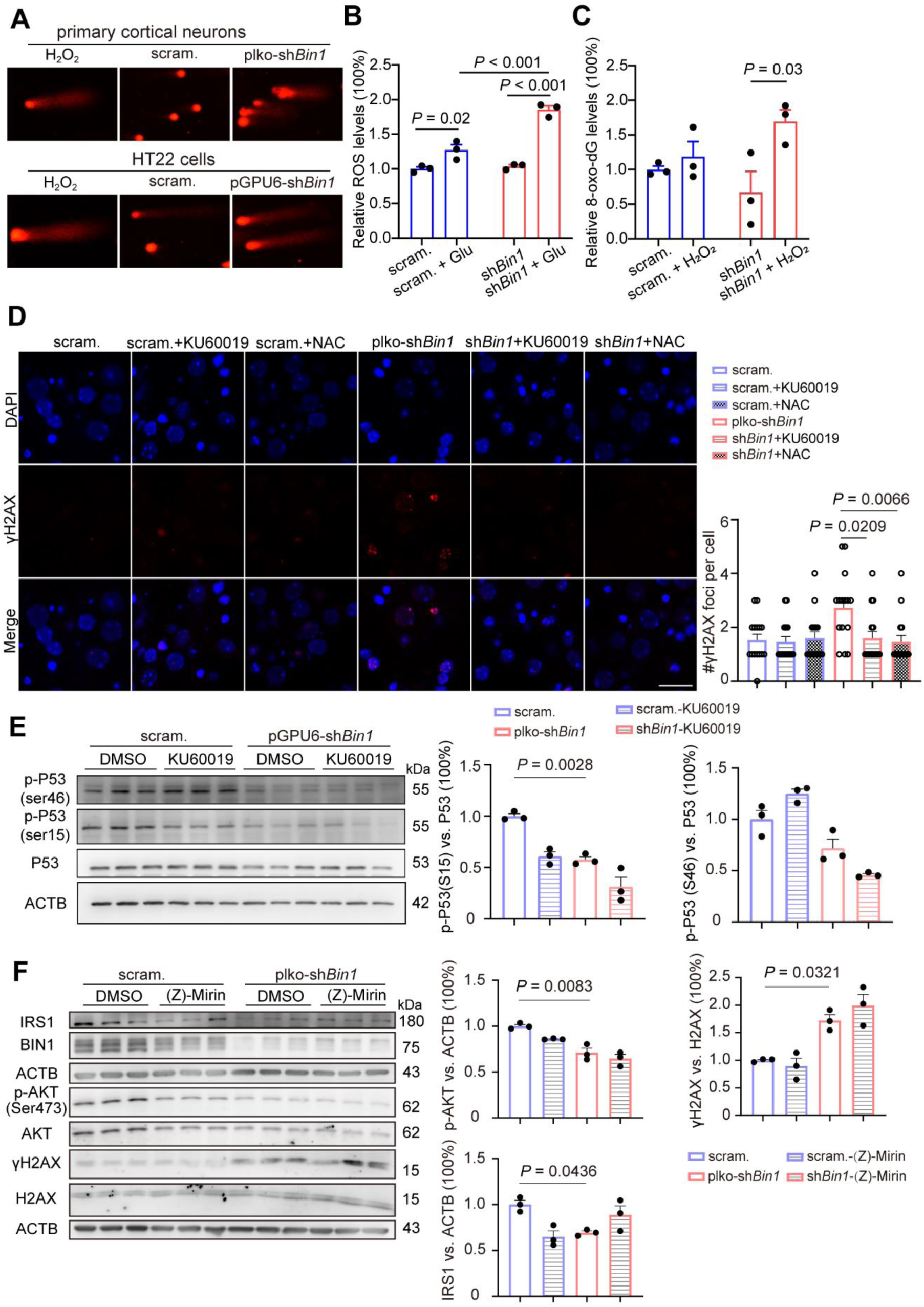
(**A**) Representative comet assay images show the DNA damage in the different groups of primary hippocampal neurons and HT22 cells. (**B**) A bar graph (Mean + SEM) overlaid with dot plots (individual data points) showing the ROS levels in scramble and pGPU6-sh*Bin1* groups with or without Glutamate treatment (5 μM, 24 h); n = 3 per group. (**C**) A bar graph (Mean + SEM) overlaid with dot plots (individual data points) showing the 8-oxo-dG levels in scramble and pGPU6-sh*Bin1* groups with or without H_2_O_2_ treatment (20 μM, 1 h); n = 3 per group. (**D**) Representative immunofluorescence images showing the γH2AX^+^ cells in the scramble and plko-sh*Bin1* groups with or without KU60019 (3 μM, 72 h) and NAC (3 mM, 12 h) treatment. Scale bar: 20 μm. n = 15 per group. (**E**) Representative WB images showing protein levels of p-53-Ser46, p-53-Ser15, and P53 in scramble and pGPU6-sh*Bin1* groups with or without KU60019 treatment (3 μM, 72 h); n = 3 per group. (**F**) Representative WB images and a bar graph (Mean + SEM) overlaid with dot plots (individual data points) showing IRS1, p-AKT-Ser473, AKT, γH2AX, H2AX, and BIN1 protein expression in scramble and plko-sh*Bin1* groups with or without Z-Mirin treatment (50 μM, 6 h); n = 3 per group.

While γH2AX foci are a gold standard marker of DSBs in cells following exposure to γ-radiation, whether they indicate genuine DSBs in ROS-stressed cells is in debate. While even biologically relevant levels of oxidative stress can induce clustered DNA lesions that subsequently develop into DSBs,^45^ treating muscle cells with SSB-inducing agents also leads to ATM-dependent γH2AX formation and γH2AX foci, which are not accompanied by p53 phosphorylation.^46^ This is interesting as p53 is a canonical substrate of ATM in DSB-induced DDR. We thus examined the phosphorylation of p53. In HT22 cells, neither p-Ser15-p53 nor p-Ser46-p53 levels are elevated in the sh*Bin1* group; in fact, p-Ser15-p53 levels are significantly decreased, and p-Ser46-p53 also displays a trend of reduction in the RNAi group (**Figure 6E**). Next, we treated the primary neurons with Mirin, a potent MRN inhibitor, to see whether it works analog to ATM inhibition; we found Mirin did not affect γH2AX levels, nor did it alter the insulin signaling (**Figure 6F**), demonstrating that BIN1 deficiency-induced ATM activation does not involve the MRN recruitment step. These results support that BIN1 deficiency does not generate genuine DSBs. A recent study identified the base excision repair (BER) endonuclease AP-endonuclease 1 (APE1) as a direct ATM activator, ^47^ providing a molecular link between SSBs and ATM. Together, we believe that BIN1 deficiency leads to SSBs-induced ATM activation.

### Alpha Lipoic Acid improves spatial cognitive performance in AAV-shBin1 mice

ATM helps to maintain cellular redox homeostasis largely through enhancing the pentose phosphate pathway (PPP), which is a major source of the reduced form of nicotinamide adenine dinucleotide phosphate (NADPH). ATM archives this via increasing expression of the rate-limiting enzyme G6PD,^48^ as well as stimulating its activity through promoting Hsp27 binding to and phosphorylating G6PD.^49^ Following neuronal *Bin1* knockdown, we observed a significant decrease in G6PD proteins (**Figure 7A**) as well as its transcript *G6pdx* (**Figure 7B**). In agreement with the reduced G6PD levels, ELISA assays revealed markedly lower NADPH content in plko*-*sh*Bin1* neurons (**Figure 7C**). The canonical Wnt/β-catenin signaling pathway regulates both glycolysis and PPP through promoting glucose uptake. Lithium chloride (LiCl) is known to enhance the Wnt/β-catenin signaling through inhibiting GSK-3β-mediated β-catenin destruction. Treating HT22 cells with LiCl (20 mM, 24 h), we found that the reduced NADPH content in pGPU6-sh*Bin1* cells was effectively restored (**Figure 7D**).

**Figure 7.**
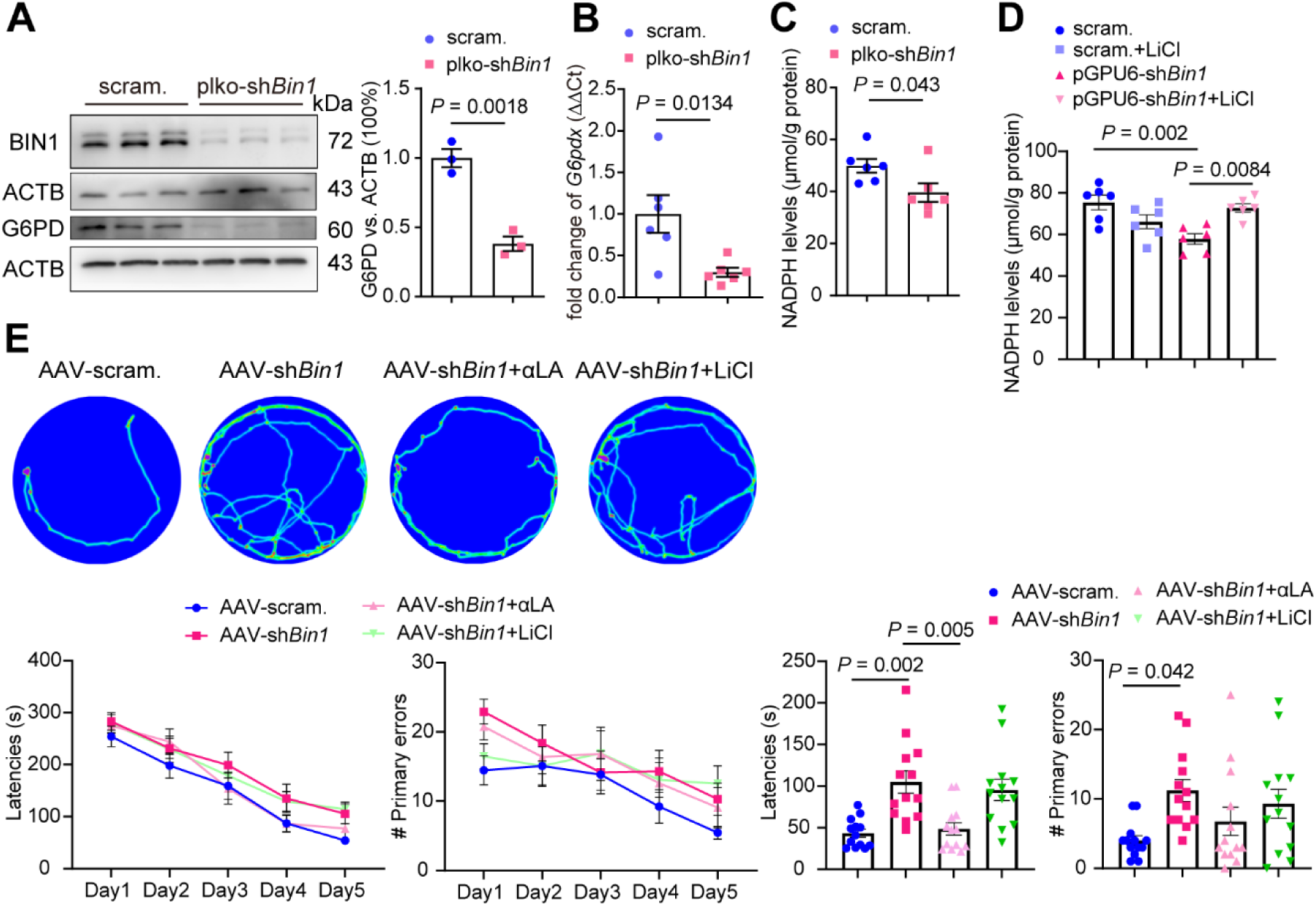
(**A**) Representative WB images and a bar graph (Mean + SEM) overlaid with dot plots (individual data points) showing G6PD and BIN1 protein expression in different groups of primary hippocampal neurons. n = 3 per group. (**B**) A bar graph (Mean + SEM) overlaid with dot plots (individual data points) showing the relative transcript levels of *G6pdx* in different groups of primary hippocampal neurons. Transcripts were examined by qRT-PCR with *Actb* as an internal reference. n = 6 per group. (**C**) A bar graph (Mean + SEM) overlaid with dot plots (individual data points) showing the NADPH levels in scramble and plko-sh*Bin1* groups; n = 6 per group. (**D**) A bar graph (Mean + SEM) overlaid with dot plots (individual data points) showing the NADPH levels in scramble and pGPU6-sh*Bin1* groups with or without LiCl treatment (20 mM, 24 h); n = 3 per group. (**E**) Heat maps showing representative trajectories of mice in the Barnes maze during the probe phase; bottom, line graphs showing the latencies and mistakes made (primary errors) before reaching the escape tunnel during the acquisition training phase; bar graphs (Mean + SEM) overlaid with dot plots (individual data points) showing the latencies and primary errors before reaching the previous escape tunnel location during the probe phase. n = 13 mice per group.

Despite the effectiveness of ATM inhibition in preserving insulin response in BIN1-deficient cellular models, ATM itself is not a suitable target at the organism level because of its vital physiological roles, and ATM insufficiency in actuality promotes systemic insulin resistance. Instead, targeting the upstream DNA damage inducer is more rational. To this end, we aimed to eliminate excessive ROS with the Wnt/β-catenin activator LiCl or by applying a direct antioxidant, the alpha lipoic acid (α-LA). As described above, one month after injection of the AAVs, mice received intraperitoneal administration of either LiCl (2 mmol/kg) or α-LA (10 mg/kg) daily for four weeks. We then performed the Barnes Maze to evaluate their spatial cognitive ability. In a typical setting, α-LA but not LiCl treatment effectively reduced the latency reaching the target holes (**Figure 7E**); the number of incorrect exploration attempts also displays a trend of reduction, but there is no significant difference (**Figure 7E**). Since both latency and exploration errors still appeared to be decreasing at the end of the training phase, we initially suspected that mice were insufficiently trained. However, retraining did not produce an appreciable additional improvement (**Figure S4**). Together, these results support the effectiveness of the anti-ROS strategy with α-LA, and the failure of LiCl might be from its insufficient strength or a cognition-impairing effect of the Lithium through other mechanisms.

### BIN1-high neurons are also more likely to express insulin signaling genes and neuronal activity genes

The above results draw a positive correlation between BIN1 protein levels and neuronal insulin signaling activity. This relationship is possibly conserved at the transcript level. To test this hypothesis, we imported a single-nucleus transcriptomic dataset (GSE157827) that contains an RNA-sequencing of the prefrontal cortex from AD patients as well as healthy control subjects. After clustering the cells into different groups, we further divided neurons into *BIN1-low* (cells with zero *BIN1* count) and *BIN1-high* (cells with nonzero *BIN1* count) subgroups. Evaluating the expression of ISP genes (*IRS1, TSC1, TSC2*) and *ATM*, we found that they are all relatively highly expressed in the *BIN1*-high subgroup (**Figure 8A**); meanwhile, neurons expressing ISP genes or *ATM* are more likely to fall into the *BIN1-high* subgroup (**Figure 8A**). Similar to the ISP genes, we also analyzed four activity-related genes (*ARC*, *BDNF*, *EGR1*, and *NR4A1*) specifically in excitatory neurons, with the results revealing a similar conserved relationship between *BIN1* and them (**Figure 8B**). *FOS* is excluded because of its more transient nature as an activity gene. It is worth noting that neuronal *BIN1* levels are also positively related to the Base Excision Repair pathway that is responsible for SSB repair (**Figure S5**). This is possibly because the chronic genotoxic stress produced by BIN1 deficiency eventually leads to defective DNA repair capability, as reported in neurons from AD patients.^50^

**Figure 8.**
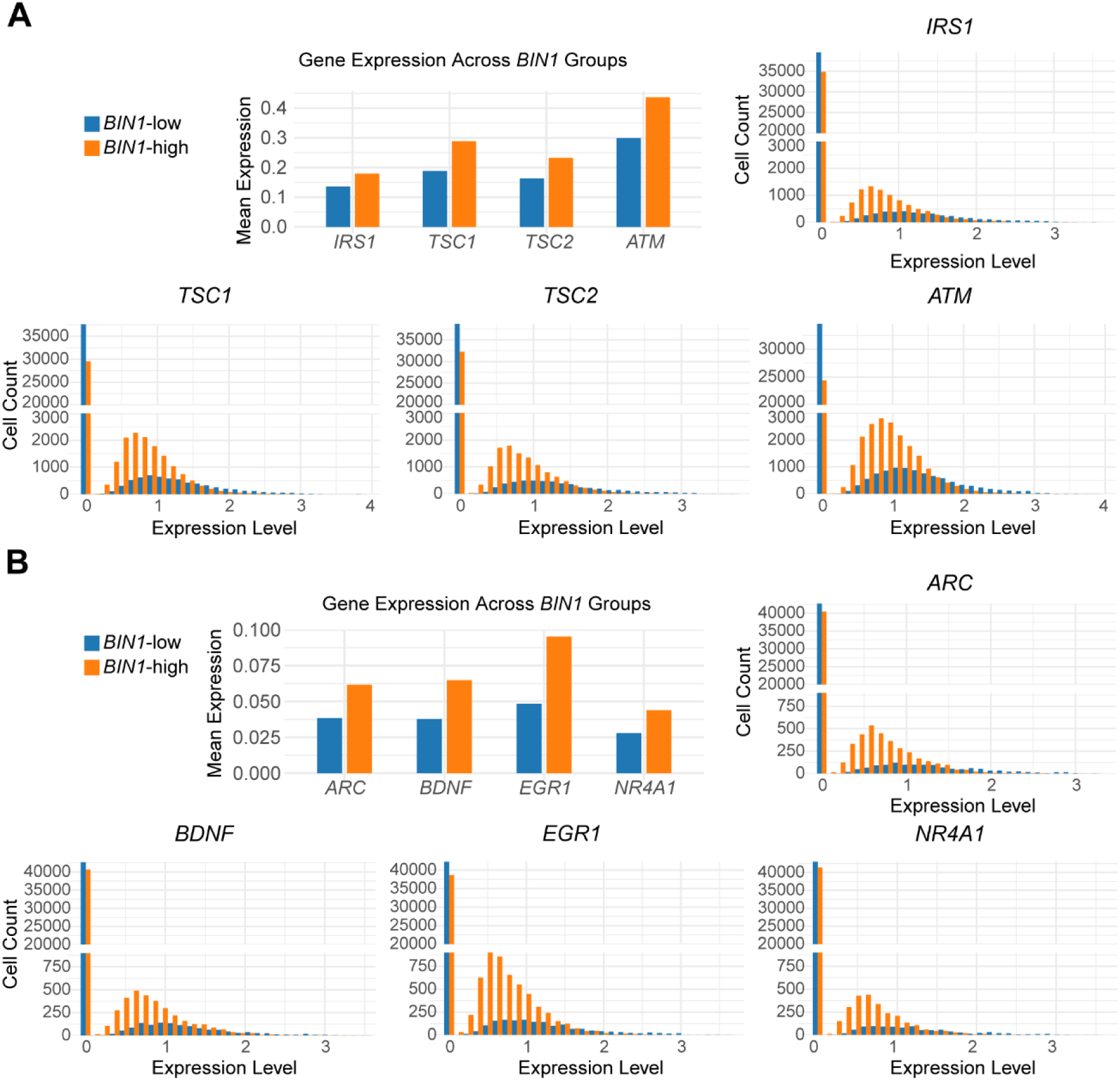
(**A**) Left, bar graph showing mean expression levels of ISP genes including *IRS1*, *TSC1*, *TSC2*, and *ATM* in *BIN1-low* (counts is zero) and *BIN1-high* (counts is non-zero) groups; right, histograms showing distribution of cell counts across the expression levels of the corresponding genes (log2 normalized counts for *IRS1*, *TSC1*, *TSC2*, and *ATM* respectively in each cell). (**B**) similar to (**A**) but plotting neuronal activity genes including *ARC*, *BDNF*, *EGR1*, and *NR4A1*.

## Discussion

In the present work, we provide evidence to propose ATM as a key mediator downstream of neuronal BIN1 deficiency. On one side, BIN1 deficiency leads to activation of nuclear ATM and subsequently ATM-dependent DDR, which decreases several key ISP proteins. The surprising effectiveness of the ATM inhibitor supports that DDR is the main route to cellular-level insulin resistance. DDR and the subsequent DNA damage repair pathways modify chromosome accessibility for DNA-binding proteins, including transcription factors, leading to large-scale alterations of gene expression.^51^ It is reported that this chromosome modification operates in a non-stochastic way, and preferentially activates stress-response transcriptional programs;^52^ but the detailed molecular mechanism is currently unclear and awaits future investigations. On the other side, nuclear ATM-mediated DDR is accompanied by a significant decrease in cytoplasmic ATM, which also contributes to insulin resistance by directly reducing AKT phosphorylation. To reconcile this seeming conflict, we propose that a small fraction of the total ATM is sufficient to support the DDR. A similar scenario was captured by researchers before; an interesting study performed by Jiali et. al found that *ATM^KO^* mice express a new catalytically active ATM protein through alternative splicing, and it effectively responds to DNA damage despite the low protein level.^53^ We also investigated the crosstalk between cytoplasmic ATM reduction and nuclear DDR. We found that cytoplasmic ATM deficiency-induced ROS hypersensitivity predisposes to oxidative DNA damage and the subsequent nuclear ATM-mediated DDR. It thus appears that autophagic degradation of the cytoplasmic ATM is a more upstream event, but it needs to be noted that the possibility of ROS-independent ATM activation is not completely excluded.

As a nexus of this ATM-centered regulation, insulin resistance is a natural interfering target and has been proven to be effective in both cellular and animal models. Neuronal insulin resistance is not new to AD researchers, and it is usually interpreted as a consequence of extracellular Aβ exposure, through binding and reducing membrane insulin receptors, or because of inflammatory factors such as TNFα and IL-1β from activated glial cells. In the present work, we report a new cell-autonomous mechanism in which DDR operates as a main driver through multi-layers regulation of the IRS1–PI3K– AKT pathway. Though DDR was known to affect the insulin signaling, previous studies focused on activated ATM as a PI3K kinase and the inhibition of PI3K–AKT signaling by its downstream effector p53. The possibility of a global change in ISP protein levels was not sufficiently recognized before. Insulin signaling promotes catabolic metabolism to support cell growth, which is an energy-intensive process; on the other hand, DNA damage repair is also highly ATP-consuming and requires the cell to relocate energy from protein/lipid synthesis to it. Meanwhile, as DDR globally affects the cell state, it requires a different set of signals from the ones directing cellular growth, additionally raising the necessity to temporally suppress the insulin signaling. Considering the prime role of genome integrity in the cellular architecture, when DNA damage happens, it is rational to reduce the response to external signals, including insulin, and orchestrate signaling pathways to support DDR and the damage repair. The above argument suggests an overall positive correlation between DDR and insulin resistance, but whether this universally applies to other cells and/or in other settings requires future investigation, considering both DDR and insulin resistance are inherently heterogeneous.

Clinical and preclinical studies have drawn a close relationship between neuronal insulin resistance and cognitive impairment. Upon binding of insulin to its receptor, the PI3K–-AKT–mTORC1 arm is activated to support those protein synthesis-required neuronal plasticity, with late-phase LTP (L-LTP) and metabotropic glutamate receptor-dependent long-term depression (mGluR-dependent LTD) in the hippocampus as two most well-known examples.^54, 55^ It is worth noting that the proper response to external signals other than a straight-lining activation of the relevant signaling pathway is critical for neuronal plasticity. In patients with loss-of-function mutations of *TSC1/TSC2*, the persistent activation of mTORC1 in fact also leads to cognitive impairment, in which specific mTORC1 inhibition is a promising therapeutic strategy under intense investigation.^56^ The other RAS–ERK1/2 arm also support neuronal plasticity especially for the above-mentioned mGluR-dependent LTD,^57^ which requires ERK activation not only because ERK1/2 and AKT–mTORC1 converge on eIF4E to regulate protein synthesis, but also because ERK1/2 phosphorylates ELK1 and the transcription factor CREB that are important for the expression of neuronal plasticity genes. The pan-mTOR inhibitor rapamycin mainly attenuates the PI3K–AKT–AKT–mTORC1 arm, and its effectiveness in our behavior experiments supports the critical involvement of this arm. For liraglutide, except for its insulin secretion-promoting effect in the peripheral, it also directly binds to neuronal GLP-1 receptors, leading to increased intracellular cAMP levels that further activate the PKA–CREB signaling, thus supporting the cognitive improvement. Therefore, while liraglutide treatment did improve p-AKT levels in AAV-sh*Bin1* mice, its cognitive improvement effect should not be interpreted as only from increased insulin sensitivity. If targeting insulin resistance is the main focus, one may also consider peroxisome proliferator-activated receptor γ (PPARγ) agonists that are represented by pioglitazone. These drugs are known to overcome insulin resistance at the cellular level, but they typically penetrate the blood-brain barrier (BBB) poorly;^58^ what’s more, though they attenuate cognitive decline in animal models,^59^ this effect disappears in AD clinical trials.^60, 61^ Leriglitazone is a recently developed brain penetrant PPARγ agonist,^62^ and future studies may explore whether it suited neurodegenerative disorders with intact BBB better. Interestingly, a previous study reported that the GLP-1 agonist increases PPARγ expression in endothelial cells.^63^ It is thus possible that liraglutide works partly through activating the PPARγ signaling. This group of drugs is promising, and one of them, semaglutide, is currently under Phase III clinical evaluations for AD treatments.

Though targeting insulin resistance is rational and has been proven to be effective, two concerns drove us to look for alternative interfering strategies. First, insulin resistance is too downstream for ATM-mediated DDR, which disturbs many other cellular processes; second, insulin resistance might not be all bad, as it also lowers energy consumption and ROS production. Indeed, some researchers believe insulin resistance at the cellular level is an evolutionarily conserved adaptive physiological mechanism aiming to control ROS, though it might produce systemic adverse effects.^64–66^ Therefore, we turned to the upstream hopping to prevent DDR from the beginning and found that the antioxidants α-LA work as expected in terms of preserving cognitive ability. There is known crosstalk between ROS and insulin signaling; ROS by itself may impair insulin signaling through protein oxidation. The antioxidant α-LA is capable of increasing insulin sensitivity and is frequently prescribed for diabetes patients with peripheral neuropathy. In this sense, α-LA appears to simultaneously target the upstream DNA damage and the downstream insulin resistance. However, the half-life of an antioxidant (e.g., 30 min to 1 hour for α-LA) is typically short, which questions their suitability for confronting chronic oxidative stress; in addition, too many antioxidants may elicit reductive stress that disturbs proteostasis and also leads to insulin resistance. Therefore, the α-LA intervention in the present study should be viewed mainly as a proof-of-concept work. Further investigations are required to explore drugs that last long and more optimized interventional regimes.

It is surprising that simply reducing the expression of an AD risk gene is sufficient to induce cellular-level insulin resistance, redox dyshomeostasis, genuine DNA damage, DDR, and autophagy elevation, which are all evidenced in AD patients and animal models. This coincidence strongly supports the loss of neuronal BIN1 as an important, and probably common, molecular event in the AD disease course, most likely in the later stage. Future studies may explore the value of neuronal BIN1 as a biomarker reflecting diverse aspects of the cell state. The present work also identifies ATM as a central mediator downstream of neuronal BIN1 deficiency, and we are thus facing a situation reminiscent of the neurodegenerative disorder AT. AT has no cue, but the antioxidant therapy displays some protective effects,^67^ which is in agreement with our results. Attenuating ROS and especially ROS-induced DNA damage, alone or in combination with an intervention improving insulin sensitivity, might also be effective for AD treatment.

## Materials and Methods

### Reagents and antibodies

Leupeptin (SG2012), bafilomycin A1 (SF2730) were purchased from Beyotime Biotechnology. Chloroquine was purchased from Beijing Vokai Biotechnology (A31639) and Macklin (C843545). SU6668 (N872395), Glutamate (#L810495) and metformin (#M813341) were purchased from Macklin. Rapamycin (GC15031), liraglutide (GC10311), and insulin (GP20673) were purchased from GlpBio. PF-4708671 (HY-15773), Hydroxyzine (HY-B0548A), KU60019 (HY-12061), LMK-235 (HY-18998), Z-Mirin (HY-117693) and NAC(HY-B0215), Cycloheximide (HY-12320) were purchased from MedChemExpress. Lithium chloride (#A100416-0025) was purchased from Sangon Biotech.

The primary antibodies used for WB and immunofluorescence were: anti-BIN1 (Proteintech, 14647-1-AP), anti-ACTB (Proteintech, HRP-60008), anti-MTOR (7C10) (Cell Signaling Technology, 2983), anti-phospho-p70S6 Kinase (Thr389) (108D2) (Cell Signaling Technology, 9234), anti-p70S6 Kinase (Cell Signaling Technology, 9202), anti-AKT (Proteintech, 10176-2-AP), anti-phospho-AKT (S473) (Proteintech, 66444-1-lg), anti-phospho-AKT (T308) (Santa Cruz, sc-271966), anti-TSC2 (Proteintech, 24601-1-AP), anti-phospho-TSC1 (S511) (Proteintech, 80340-1-RR), anti-TSC1 (Proteintech, 29906-1-AP), anti-γH2AX (Proteintech, 29380-1-AP), anti-H2AX (Proteintech, 10856-1-AP), anti-Tau (Proteintech, 10274-1-AP); anti-phospho-Tau (Ser396) (abcam, ab32057), anti-phospho-ATM (S1981) (ProMab, P30249); anti-ATM (Abways, CY7229); anti-c-Fos (Affinity, AF0132); anti-G6PD (Proteintech, 66373-1-Ig); anti-EIF4EBP1 (Beyotime, AF5159); anti-phospho-EIF4EBP1 (Beyotime; AF5806). The secondary antibodies used for WB and immunofluorescence were: HRP-conjugated Goat anti-Mouse IgG (Proteintech, SA00001-1), HRP-conjugated Goat anti-Rabbit IgG (Proteintech, SA00001-2), CoraLite488-conjugated Goat anti-Mouse IgG (Proteintech, SA00013-1), CoraLite 488-conjugated Goat anti-Rabbit IgG (Proteintech, SA00013-2), Rhodamine (TRITC)-conjugated Goat anti-Mouse IgG (Proteintech, SA00007-1), Rhodamine (TRITC)-conjugated Goat anti-Rabbit IgG (Proteintech, SA00007-2), and Alexa Fluor™ Plus 405-conjugated Goat anti-Rabbit IgG (Invitrogen, A48254).

### RNA interference, lentivirus, and AAVs

The siRNAs were all synthesized by Genepharma with sequences shown below: si*Bin1*: 5’-GCTCAATCAGAACCTCAATGAT-3’; si*Ulk3*: 5’-GGUUAUUUCUAAAGUUAGA-3’; To reduce *Bin1* expression in cultured primary neurons, we constructed a lentivirus vector expressing an shRNA targeting all isoforms of mouse *Bin1* under the Rnu6 promoter as well as an EGFP reporter under a separate *hPGK1* promoter for convenient visualization of successfully transduced cells under a microscope (plko-U6-sh*Bin1*-hPGK-EGFP); the control virus expresses a scrambled sequence of sh*Bin1* (plko-U6-scramble-hPGK-EGFP). Both lentiviruses were packaged and produced by Hanbio.

To reduce neuronal *Bin1* expression in the hippocampal CA1, we employed a rAAV2/9 *Mir30a*-based vector expressing the same shRNA targeting all isoforms of mouse *Bin1*, as well as a *mCherry* reporter, both under the *Camk2a* promoter for specifical expression in excitatory neurons (rAAV-*Camk2a*-mCherry-5’*MiR30a*-sh*BIN1*-3’*MiR30a*-WPREs, abbreviated as AAV-*Camk2a*-*mCherry*-sh*Bin1* or AAV-sh*Bin1*); rAAV-*Camk2a*-mCherry-5’*MiR30a*-scramble-3’*MiR30a*-WPREs (abbreviated as AAV-*Camk2a*-*mCherry*-scramble or AAV-scram.) was used as the control. The two AAVs were packaged by BrainVTA.

### Culture of primary hippocampal neurons

The primary hippocampal and cortical neurons were isolated from newborn (P0) mice with a standard procedure reported before ^68^. Briefly, mice were disinfected with 75% alcohol before sacrifice and dissection; in cold DMEM (Thermo Scientific, C11995500BT), the meninges were carefully and gently peeled off with forceps, and the hippocampus and the cortex were identified and dissected out; the hippocampus and cortex were then cut up and digested in a solution containing 1 mg/ml DNase (Merck, D4513) and 2 mg/ml papain (Sangon Biotech, 9001-73-4), filtered through a Falcon® 40 µm cell strainer (Corning, 352340); cells were then transferred to Neurobasal™-A (Thermo Scientific, 10888022) containing 2% B-27™ (Thermo Scientific, 17504044), 1% penicillin/streptomycin (Thermo Scientific, 15140122), and 2 mM L-glutamine (Thermo Scientific, 35050061). Neurons were cultured in Dishes or plates coated with 0.01 mg/ml poly-D-lysine (Merck, P6407) for 5 days before transfection or lentivirus transduction.

### Lentivirus transduction

The lentivirus was introduced into the primary hippocampal neurons at 5 days *in vitro* (DIV5). 24 h after the transduction, the old medium containing the virus was removed and replaced with fresh medium. On DIV11, neurons were harvested for the subsequent immunofluorescence, real-time quantitative reverse transcription PCR (qRT-PCR), or WB analyses.

### Mouse maintenance, separation of brain tissue

The 8-week-old wild-type C57BL/6J mice were purchased from the Animal Core Facility of Nanjing Medical University. All mice underwent a 12-hour light/dark cycle under appropriate ambient temperature and humidity, with good ventilation, and ad libitum access to food and water. Mice were injected with AAV-*Camk2a*-*mCherry*-sh*Bin1* or AAV-*Camk2a*-*mCherry*-scramble in bilateral hippocampal CA1 and fed for 8 weeks before the sacrifice. Mice were anesthetized by intraperitoneal injection of 30 ml/kg phenobarbital (Merck, P3761). After perfusion with cold physiological saline, the skull was opened, and the whole brain tissue was collected. All tissues were frozen in liquid nitrogen and stored at −80℃ before use.

### Stereotaxic intracranial injection

Mice were anesthetized and fixed in a stereotaxic frame; the hair near the injection site was cut off and disinfected twice with betadine. Carefully cut the skin along the sagittal slit with disinfected scissors to expose the skull surface. For the introduction of AAVs into the bilateral hippocampal CA1, 0.5 µL of virus for each side was injected at the following coordinates: 2.5 mm behind bregma, 1.0 mm off, and 1.9 mm deep. The infusion rate was 0.1 µL /min with a Hamilton 33G syringe needle, which remained at the injection site for 5 minutes after the infusion to prevent virus spillover. After the injection, mice were placed in an isolated environment to facilitate their recovery. All experiments were performed 8-10 weeks after the injection.

### Intraperitoneal injection

Mice were intraperitoneally injected with 2 mg/kg rapamycin or 200 µg/kg liraglutide for 4 weeks after AAV infection for 8-10 weeks. The control group received an equal amount of PBS by intraperitoneal injection.

### Behavioral tests

Mice’s behavioral images were captured with a computer-connected digital camera (TopScan CleverSys, USA). The test mice were acclimated to the room for 2 hours before a behavioral test. Learning and memory abilities were assessed with the Barnes Maze test.

### Barnes maze

Barnes maze is used to evaluate long-term spatial memory in mice ^69^. The behavioral test consists of five days of training and one day of testing. The experimental device is a disk with a diameter of 90 cm. There are 22 holes evenly distributed at 5 cm from the edge of the disk, of which only one hole is equipped with an escape box. A bright light source was placed above the platform, forcing the mice to explore and hide in the box. During the training period of 5 consecutive days, the mice were trained once a day. If the mice did not enter the escape box within 300 s, they were guided into it, and the mice were removed and put back into the cage 30 s later. The tests were conducted three days apart after the training phase. During the test period, each mouse was active for 300 s. Software was used to record data in real time during behavioral processes. Finally, the time when mice entered the escape box and the number of times they sniffed the wrong hole were counted.

### ROS detection

ROS levels were evaluated with a DCFHDA-based ROS detection kit (Beyotime, S0033S) according to the manufacturer’s manual. Briefly, after removing the cell culture medium, an appropriate volume of diluted DCFHDA was added, and cells were then incubated in the incubator at 37℃ for 20 min. Cells were washed three times with serum-free culture medium to fully remove the DCFHDA that did not enter cells before digestion with 0.25% pancreatin. Cells were then collected and detected with a Varioskan Flash microplate reader (Thermo Scientific, USA). The excitation wavelength of 488 nm and the emission wavelength of 525 nm were used for the detection.

### NADPH detection

The kit (No. BYSH-0208W) was purchased from Boyan Biotechnology. According to the instructions, NADPH was extracted, and a given amount of extract B was used to lyse the cells. The lysed cells were collected into an EP tube using a cell scraper on ice at 4℃ for 30 min. After incubation, they were immediately placed on ice for 5 min. The samples were then transferred to a new EP tube after 4℃ pre-cooling centrifugation, with a speed of 12000 rpm for 10 min. Half of the supernatant was aspirated and transferred to a new EP tube. Extract A was added in small amounts multiple times, and the solution was tested with a pH paper for color development. The pH of the resulting solution was adjusted to neutral, and the volume of extract A added to each sample was recorded.

Under 4℃ conditions, the samples were centrifuged at 12000 rpm for 5 min, and the supernatant was transferred to a new EP tube and placed in an ice bath before testing. According to the instructions, reagents one, two, and three (powders should be centrifuged before dissolution in distilled water) were added to the 96-well plate in the corresponding amounts. The samples were incubated in the dark at 37℃ for 10 min, followed by the addition of reagent four. The microplate reader was set to 37℃ and 450 nm wavelength, and the absorbance values of each well were measured and recorded as A1. Thirty minutes later, the absorbance values of each well were measured again using the same parameters on the microplate reader and recorded as A2. The final content was calculated using the formula provided in the manual.

### 8-oxo-dG detection

#### Pre-treatment of the samples and ELISA

On DIV11, neurons were exposed to 20 μM H_2_O_2_ for 1 h, and then collected in PBS. Cellular DNA was extracted through the freeze-thaw cycle: samples were frozen to −80°C and then immediately transferred to a 37°C water bath for thawing for a total of three cycles. Samples were then centrifuged at 12000 rpm at 4°C for 15 mins, with the supernatant collected for the subsequent ELISA steps and the precipitate for protein extraction.

For ELISA, the mouse 8-oxo-dG ELISA Kit (Huabo Deyi, Beijing, China) was employed. Samples were analyzed in triplicate (50 µL sample/well) according to the manufacturer’s instructions. Concentration of the 8-oxo-dG (ng/ml) was then normalized to the protein concentration (nmol 8-oxo-dG/mmol protein).

### Cell line culture and transfection

Immortalized mouse hippocampal HT22 cells (Wuhan Pricella Biotechnology, CL-0697) were cultured in DMEM high-glucose medium containing 10% fetal bovine serum with penicillin and streptomycin. Transfections were performed with Lipofectamine™ 3000 (Invitrogen, L3000075) when cells reached ∼70% confluence. Transfected cells were cultured in the incubator (37℃, 5% CO_2_) for an additional 72 h before harvest.

#### Comet electrophoresis

Also known as single-cell gel electrophoresis, the collected cell suspension is evenly spread on a microscope slide previously coated with 1% normal melting point agarose in a 37℃, 75 μL, 0.7% low-melting-point agarose solution, and incubated at 4°C for 10 min to solidify. The slide is then incubated in the lysis solution at 4℃, 1-2 h. After lysis, the slide is incubated at room temperature in an alkaline electrophoresis buffer for 20-60 min to unwind the DNA, followed by electrophoresis (25 V) for 20-30 min. The slide is then placed in a neutralizing buffer (0.4 M Tris, pH 7.5) at 4°C for 5-10 min, after which the buffer is discarded and the slide is incubated in the dark with 20 μL Propidium Iodide Solution for 15 min. The slide is washed three times with deionized water, covered with a coverslip, and finally observed under a fluorescence microscope to obtain images.

#### ELISA

Collect the supernatants from primary neuronal cultures of each group. According to the instructions for the mouse Aβ_1-42_ and Aβ_1-40_ ELISA kit, add the prepared supernatants and standards to the reaction wells, incubate at 37℃ for 30 min. After incubation, wash the wells 5 times, then add the enzyme-labeled reagent and react at 37℃ for another 30 min. Following the washing steps, add the chromogenic solution and allow a 10-minute color development reaction. Finally, add the stop solution to terminate the reaction, and read the absorbance values of each well within 15 min using an enzyme-labeled microplate reader.

### Immunofluorescence

Tissues were fixed with 4% paraformaldehyde for 24 h, dehydrated with 20% sucrose in Phosphate-buffered saline (PBS, Sangon Biotech, E607008) for 24 h, and then 30% sucrose in PBS for ∼24 h until they sank to the bottom. Tissues were embedded in OCT glue (Sakura Finetek, 4583), sliced at a thickness of 20 μm with a CM1950 cryostat microtome (Leica Biosystems, Germany), and lastly mounted on gelatin-coated slides. 0.1% Triton X-100 (Sangon Biotech, A600198) for 1 h was applied for tissue permeabilization, and then tissues were blocked in 5% BSA (Sangon Biotech, A600332). For cultured cells, the permeabilization time is 10 min. For primary antibodies, samples were incubated at 4℃ overnight; for secondary antibodies, samples were incubated at room temperature for 2 h.

### WB

RIPA buffer (Beyotime, P0013B) with a protease inhibitor cocktail (Beyotime, st506) and a phosphatase inhibitor cocktail (Roche, 04906837001) was used for cell lysis and protein extraction. For tissue homogenization, cells or tissue lysate were placed in an ice shaker at 4℃ for 30 min to fully lyse. With 10000 x g centrifugation, supernatant was collected and protein quantification was performed using Enhanced BCA Protein Assay Kit (Beyotime, P0010S). The SDS-PAGE Sample Loading Buffer 5X (Beyotime, P0015) was added to the remaining protein lysate, and the protein was heated at 95℃ for 5 min. Thirty-μg protein samples were separated by electrophoresis at 80 v, 30 min, 120 v, 2 h, and then transferred to Immobilon®-PSQ PVDF Membrane (Merck, ISEQ08100), blocked with 5% dry skim milk, incubated with different primary antibodies in a shaking bed at 4℃ overnight, and washed with TBST for 3 times, 10 min each time. After incubation for 1 h, the corresponding HRP-conjugated secondary antibody was washed with TBST 3 times, 10 min each time. When there are too many samples to fit in a single gel but analyzing in one batch is necessary, we run two gels and the following procedures alongside to enable a cross-comparison between groups on different gels.

### qRT-PCR

RNAiso Plus (Takara Bio, 9109) was used to extract total RNA from cells or tissues. 4x EZscript Reverse Transcription Mix II (with gDNA Remover) kit (EZBioscience, EZB-RT2GQ) and 2x SYBR Green qRT-PCR Master Mix (EZBioscience, A0001-R1) were used for qRT-PCR with the following settings: 95℃ for 5 min, 60℃ for 30 s for 40 cycles, 95℃ for 10 s; *Actb* was employed as an internal reference; the delta-delta Ct method was used to calculate the relative fold of target genes. All primers were synthesized by Sangon Biotech. Sequences of all primers were listed in **Table S1**.

### Image analysis

A DM4000 B fluorescence microscope (Leica Biosystems, Germany) and an LSM 710 laser scanning confocal microscope (Zeiss, Germany) were employed to image all cell slides or tissue sections. For each group, at least 3 slides were randomly imaged. For immunostaining or fluorescence intensity analysis, ∼3 regions of interest (ROIs) with the fixed area were randomly selected for each neuron from high-magnification images; averaged intensities of each ROI were employed for comparison.

### Single-cell transcriptomics Data Source and Analysis

The GSE157827 dataset, which is from single-nucleus RNA-sequencing of prefrontal cortex from AD patients and healthy control subjects, was re-analyzed in the present work. Briefly, after standard quality control and preprocessing, a total of 166,685 cells were identified, of which 88,363 were annotated as neurons. Neurons were then stratified into *BIN1-high* (non-zero *BIN1* expression) and *BIN1-low* (zero expression) groups based on *BIN1* transcript levels. To eliminate the influence of different group sizes, 43993 cells in each group were employed (randomly selected when necessary) for the plotting analysis. Expression of selected ISP genes (*IRS1, TSC1, TSC2, ATM*) was compared between the two groups. Data visualization was performed using grouped bar plots and histograms.

Employing the same analytical framework, we also analyzed expression of neuronal activity-related genes (*ARC, BDNF, EGR1*, and *NR4A1*) and base excision repair (BER)– related genes (*OGG1, NEIL1, NEIL2, APEX1, XRCC1, PARP1*) between *BIN1*-defined subgroups in excitatory neurons (n = 62229 in total; n = 29679 in each subgroup). Excitatory neurons were defined by simultaneous high expression of *SLC17A7* (encoding VGLUT1) and *CAMK2A* (encoding calcium/calmodulin-dependent protein kinase II alpha) in the annotated neurons.

### Study approval

All animal experiments were conducted following the international standards on animal welfare and the guidelines of the Institute for Laboratory Animal Research of Nanjing Medical University (approval no. IACUC: 2201039 and IACUC:1812054-4).

### Statistical analyses

All data were statistically analyzed by GraphPad Prism. Data were expressed as Mean + SEM unless otherwise noted. Two-tailed Student’s t-test was performed for comparing two groups of data, and two-way ANOVA followed by Tukey’s post hoc test was employed for comparisons among 4 or more groups. *P* < 0.05 was considered statistically significant.

## Data availability statement

The data used and analyzed during the current study are available from the corresponding author upon request.

